# From Colonisation to Invasion: Genomic and Phenotypic Comparison of Faecal and Bloodstream Isolates from the same patients

**DOI:** 10.1101/2025.07.15.664880

**Authors:** Aakash Khanijau, Ellie Allman, Ralfh Pulmones, Richard N. Goodman, Rachel McGalliard, Christopher M. Parry, Enitan D. Carrol, Adam P. Roberts

## Abstract

Gram-negative bloodstream infections (GNBSI) carry a significant global health burden. *Escherichia coli* and *Klebsiella pneumoniae* are the two most common causes of healthcare-associated GNBSI, which may arise from gastrointestinal tract (GIT) colonisation. Understanding genomic and phenotypic adaptations that underpin transition from GIT colonisation to invasive bloodstream infection could improve understanding of pathogenesis.

This study identified ‘linked’ faecal and blood isolates from children with healthcare-associated GNBSI caused by *E. coli* and *K. pneumoniae*. Linked pairs were compared for antimicrobial resistance, biofilm formation, and underwent comparative genomic analysis via whole-genome sequencing, comparative average nucleotide identity (ANI) and core genome single nucleotide polymorphism (SNP) analysis.

Five isolate pairs (three *E. coli*, two *K. pneumoniae*) showed high relatedness, supporting GIT origin of bloodstream infection. Isolates within pairs had identical virulence genes whereas phenotypic assays revealed changes in antimicrobial susceptibility, with one pair undergoing changes in resistance gene profiles, and increased biofilm formation in 4/5 isolates.

This study provides insight into within-host evolution from gastrointestinal colonisation to bloodstream invasion in Gram-negative pathogens. Convergence on metabolic adaptation and biofilm formation suggests these traits may be advantageous in healthcare-associated GNBSI. Further studies involving larger cohorts alongside functional validation of mutations are needed to better understand GNBSI pathogenesis.

## Introduction

Gram-negative bloodstream infections (GNBSI) contribute significantly to the global burden of paediatric sepsis, accounting for 30-50% of all bloodstream infections worldwide [1, 2]. Healthcare-associated bloodstream infections, such as central-line associated bloodstream infections (CLABSI), often affect neonates or other critically ill children, and are associated with significant mortality. [3].

The two most clinically significant organisms implicated in GNBSI are *Klebsiella pneumoniae* and *Escherichia coli* [4, 5]. The emergence of antimicrobial resistance (AMR) in these pathogens, presents a significant challenge. Particularly, extended-spectrum beta-lactamase (ESBL)-producing isolates, which exhibit resistance to third generation cephalosporins and beta-lactam/beta-lactamase inhibitor combinations. As such, the World Health Organization has designated ESBL-producing *K. pneumoniae* and *E. coli* as priority pathogens for which new antibiotics are required [6].

Both *K. pneumoniae* and *E. coli* are persistent colonisers of the human gastrointestinal tract (GIT) which is believed to be an important reservoir for subsequent GNBSI [7]. The transition from GIT colonisation to GNBSI can occur via invasion of other body sites or medical devices, leading to secondary bloodstream infection, or direct translocation across the gut mucosa causing primary bloodstream infection. The latter is implicated in 20-50% of cases of healthcare associated infections [7].

The transition from GIT colonisation to bloodstream invasion depends on host, clinical, and environmental influences, combined with specific pathogen characteristics. Host factors that can predispose to GIT-bloodstream transition include immunosuppression, critical illness, and antimicrobial exposure [7, 8]. Pathogen factors include advantageous phenotypes such as AMR, bloodstream fitness and biofilm formation; which may allow invasion, survival and dissemination in the bloodstream. Biofilm formation is of particular interest in the context of CLABSI, as growth within a biofilm provides protection against host defence and antimicrobial agents, and may act as a nidus for ongoing bloodstream dissemination.

Many antimicrobial resistance genes (ARGs) and virulence genes underpinning AMR and invasive infection respectively are well characterised. However, predisposing genotypes and specific genetic changes undergone within-host that may underpin GIT-blood transition are seldom studied, perhaps due to the scarcity of such paired isolates from the same patient. This includes genomic adaptations like acquisition or loss of virulence genes and plasmids, and regulatory or metabolic alterations. Furthermore, the relationship between many genomic adaptations and advantageous phenotypic changes remains unclear, as well as the relationship between AMR and invasive infection. Combined phenotypic and genomic comparison of faecal and bloodstream isolates, obtained from the same patient, could improve understanding of intra-host evolution involved in the pathogenesis of GNBSI, and translate to informing intervention strategies.

In this study we aimed to identify genomically ‘linked’ pairs of faecal and blood Gram-negative isolates, which were likely to have undergone GIT-blood transition and evolution within patients to cause GNBSI. For each ‘linked’ pair, we investigated if there had been any loss or acquisition of virulence genes, ARGs, and plasmids between faecal isolates and their blood counterpart. We then investigated if there were any difference in two phenotypes of interest: AMR and biofilm formation. Finally, we used a comparative genomic approach to identify further genomic changes that could explain any observed phenotypic differences or otherwise contribute to the transition from GIT colonisation to bloodstream invasion.

## Materials and Methods

### Clinical isolates, clinical data collection and ethics

A library of 81 stored bacterial isolates from Alder Hey Children’s Hospital, Liverpool, UK, was interrogated for pairs of ESBL-producing *E. coli* and *K. pneumoniae* isolates from the same patient, where the same pathogen had been identified on a routine faecal surveillance culture and in a subsequent blood culture. Isolates were stored in LB (Luria-Bertani) Broth (Sigma-Aldrich) and 20% glycerol at −80°C. Anonymised clinical information was obtained from patient records, including: demographics, source of GNBSI, and antimicrobial exposure between isolate dates. Ethical approval for linking anonymised data was granted by the University of Liverpool as part of a separate study for which the isolate library was created. (REC reference 255669).

### Whole Genome Sequencing and comparative genomics

Short-read Whole Genome Sequencing (WGS) and assembly was performed at the Centre of Genomic Research (University of Liverpool) using the Illumina NovoSeq SP (2×150 bp). Assembled genomes were analysed for the following characteristics, which were compared between faecal and blood isolates of each patient pair. Core genome multi-locus sequence typing (cgMLST) was performed using Kleborate [9] for *K. pneumoniae* isolates and using Institut de Pasteur database (https://bigsdb.pasteur.fr/about/) for *E. coli* isolates. Kaptive [10] was used to identify wzi-type, K-type and O-type of *K. pneumoniae* isolates, and SeroTyper 2.0 and CHtype to determine the H-type, O-type and FumC-type of *E. coli* isolates.

To assess relatedness further, average Nucleotide Identity (ANI) was then compared between isolates using OrthoANI [11] Following this, the number of core genome single nucleotide polymorphisms (SNPs) between faecal and blood isolates of each pair was calculated. SNPs were predicted using Snippy v.4.3.6 (https://github.com/tseemann/snippy) by aligning raw reads of assemblies to the reference genomes MG1655 and HS11286, for *E. coli* and *K. pneumoniae* respectively. Core genome SNPs were extracted with Snippy-core and intra-species SNPs were plotted and counted in R v.4.3.1 [12]. ResFinder 4.2.2 [13] was used to determine the presence of ARGs and provide a genotypic prediction of resistance for each isolate. VF analyzer (http://www.mgc.ac.cn/VFs/) was used to identify virulence genes in each isolate. PlasmidFinder [14] was used to identify the presence of plasmids.

Isolate genomes were annotated using RAST annotation tool [20] and pairs underwent comparative genomic analysis using Breseq version 0.38.3 [21] using default parameters. Mutations between blood isolates and the putative counterpart faecal ancestor were categorized as insertions/deletions (indels), non-synonymous SNPs (nSNPs), synonymous SNPs (sSNPs), or intergenic SNPs (iSNPs). Mutations within or <100bp upstream of a gene were classified into COG functional categories using EGGNOG 5.0. Mutations involving hypothetical proteins were classified as ‘Function unknown’ and mutations without genes within 100bp were ‘Unclassified.’ Any mutations relating to virulence genes, ARGs, or previously described bloodstream survival factors were identified. In addition, mutations in genes previously reported to affect antimicrobial resistance or biofilm formation were identified.

### Data access

The bioproject number for this study is PRJNA1295786 which contains the 10 accession numbers for the individual genomes SRX29836134 - SRX29836143.

### Antimicrobial Susceptibility Testing (AST)

Six antimicrobials from the contemporaneous local prescribing guidance for Gram-negative infections were used for AST: Cefotaxime, Gentamicin, Ciprofloxacin, Amoxicillin-Clavulanic acid (Co-amoxiclav), Piperacillin-Tazobactam and Meropenem. Kirby-Bauer disc diffusion testing was used for initial AST following EUCAST guidelines [15]. Zone sizes were interpreted as per the EUCAST clinical breakpoints, to give a phenotype of Sensitive (S), resistant (R) or in the ‘Area of Technical Uncertainty’ (ATU) [16]. Three technical and three biological repeats were performed, and the mode was recorded as the phenotype for each isolate, and compared within each pair. In addition, the mean zone size, along with standard error of the mean (SEM) was recorded for each isolate and Mann-Whitney U test was used to compare differences between faecal and blood isolates for each patient pair. Any identified differences between faecal and blood isolates in each patient pair were confirmed using broth microdilution to quantify the fold-change in minimum inhibitory concentration (MIC). Broth microdilution was performed following EUCAST guidelines [17] and with three biological and three technical replicates.

### Biofilm formation

Crystal violet assay was performed derived from two previously published protocols [18, 19]. Isolates were incubated at 37°C on LB agar for 18 h. A single colony of each isolate was transferred to 10ml Mueller-Hinton broth 2 (MHB2; Sigma-Aldrich) and incubated at 37°C at 200 rpm for a further 18 h. Cultures were diluted to an OD_600_ of 0.01 in fresh MHB2. 150µl of each isolate was transferred in a U-bottom plate, with 150µl sterile MHB2 used as a negative control. After 24-hour incubation, the biomass was transferred to a flat-well plate, which was used to measure OD_600_. The remaining biofilm in the U-bottom plate wells was stained with 1% crystal violet and subsequently solubilized with 30% acetic acid. The final OD_600_ was measured in a flat-well plate. This value was then normalized to the initial OD_600_ of the biomass, to quantify biofilm formation. The assay was performed as three biological replicates and four technical replicates. Mean OD_600_ of blood and faecal isolates were compared using a paired t-test, and SEM calculated.

### Statistical analysis and data visualization

All statistical analysis and data visualization was performed using ggplot2 in R version 4.3.2 [12].

## Results

### Clinical isolates and clinical context of GNBSI

Five pairs of isolates, two *K. pneumoniae* and three *E. coli,* were identified for use in this study (Table 1). Time between faecal (1st sample) and blood sample isolation ranged from 17-144 days. Patients from whom isolates were derived were aged 1 month to 12 years. All five cases of GNBSI were diagnosed as CLABSI, and 4/5 cases occurred on ICU. Patients received between two to six different antimicrobials in-between faecal and blood sample isolation.

**Table 1:** Characteristics of isolate pairs and clinical data of patients with GNBSI.

| Pair | Species of isolate pair | Days from faeces - blood isolation | Age at GNBSI | CLABSI | ICU patient at time of GNBSI | Antibiotic exposure between faeces and blood isolate |
| --- | --- | --- | --- | --- | --- | --- |
| 1 | *K. pneumoniae* | 144 | 12 years | Y | N | Ciprofloxacin, Clindamycin, Ceftriaxone, Piperacillin-Tazobactam, Co-trimoxazole, Gentamicin |
| 2 | *K. pneumoniae* | 87 | 11 months | Y | Y | Ciprofloxacin, Cefalexin, Piperacillin-Tazobactam, Ceftriaxone, Vancomycin, Gentamicin |
| 3 | *E. coli* | 185 | 9 months | Y | Y | Co-amoxiclav, Cefalexin |
| 4 | *E. coli* | 31 | 12 months | Y | Y | Ciprofloxacin, Meropenem |
| 5 | *E. coli* | 17 | 1 month | Y | Y | Ciprofloxacin, Vancomycin |

### Genomic relatedness of faecal-blood isolate pairs

To initially assess relatedness between faecal and blood isolates, cgMLST was performed. In 5/5 pairs, faecal and blood isolates shared the same sequence type (Table 2.) In Pair 1, *K. pneumoniae* isolates were from the sequence type ST922, and in Pair 2 *K. pneumoniae* isolates were from the sequence type ST461 (Table 2A). In Pair 3 and 4, *E. coli* isolates were from the globally dominant sequence type ST131, whereas in Pair 5 isolates were from sequence type 12 (Table 2B). *K. pneumoniae* isolates were then compared for K-type, O-type and wzi-type, which were concordant between faecal and blood isolates in both pairs. *E. coli* isolates were compared or O-type, H-type and FumC -type, and again, all were concordant within each pair.

**Table 2A:** MLST, K-type, O-antigen type and wzi-type of *K. pneumoniae* isolates within pairs, determined using Kleborate and Kaptive.

| Species | *Klebsiella pneumoniae* |  |  |  |
| --- | --- | --- | --- | --- |
| Pair | Pair 1 |  | Pair 2 |  |
| Isolate | Faeces | Blood | Faeces | Blood |
| MLST | ST922 | ST922 | ST461 | ST461 |
| K type | K28 | K28 | K25 | K25 |
| O type | O1 | O1 | O1 | O1 |
| wzi type | wzi84 | wzi84 | wzi25 | wzi25 |

**Table 2B:** Phylotype and MLST of *E. coli* isolates within pairs using ClermonTyping and Institut de Pasteur database.

| Species | *Escherichia coli* |  |  |  |  |  |
| --- | --- | --- | --- | --- | --- | --- |
| Pair | Pair 3 |  | Pair 4 |  | Pair 5 |  |
| Isolate | Faeces | Blood | Faeces | Blood | Faeces | Blood |
| MLST | ST131 | ST131 | ST131 | ST131 | ST12 | ST12 |
| FumC type | C40 | C40 | C40 | C40 | C13 | C13 |
| O type | H4 | H4 | H4 | H4 | H1 | H1 |
| H type | O25 | O25 | O25 | O25 | O4 | O4 |

Next, relatedness was further assessed using ANI (Figure 1A.) In 4/5 pairs (2, 3, 4, 5) the faecal-blood isolates shared an ANI of 99.99%, whereas isolates in Pair 1 shared an ANI of 99.82%. Within a comparison between all isolates across all patient pairs, each faecal isolate was most closely related to its corresponding blood isolate from the same patient. Given these findings, all pairs were at this point considered to be genomically ‘linked’ and included for further study.

**Figure 1:**
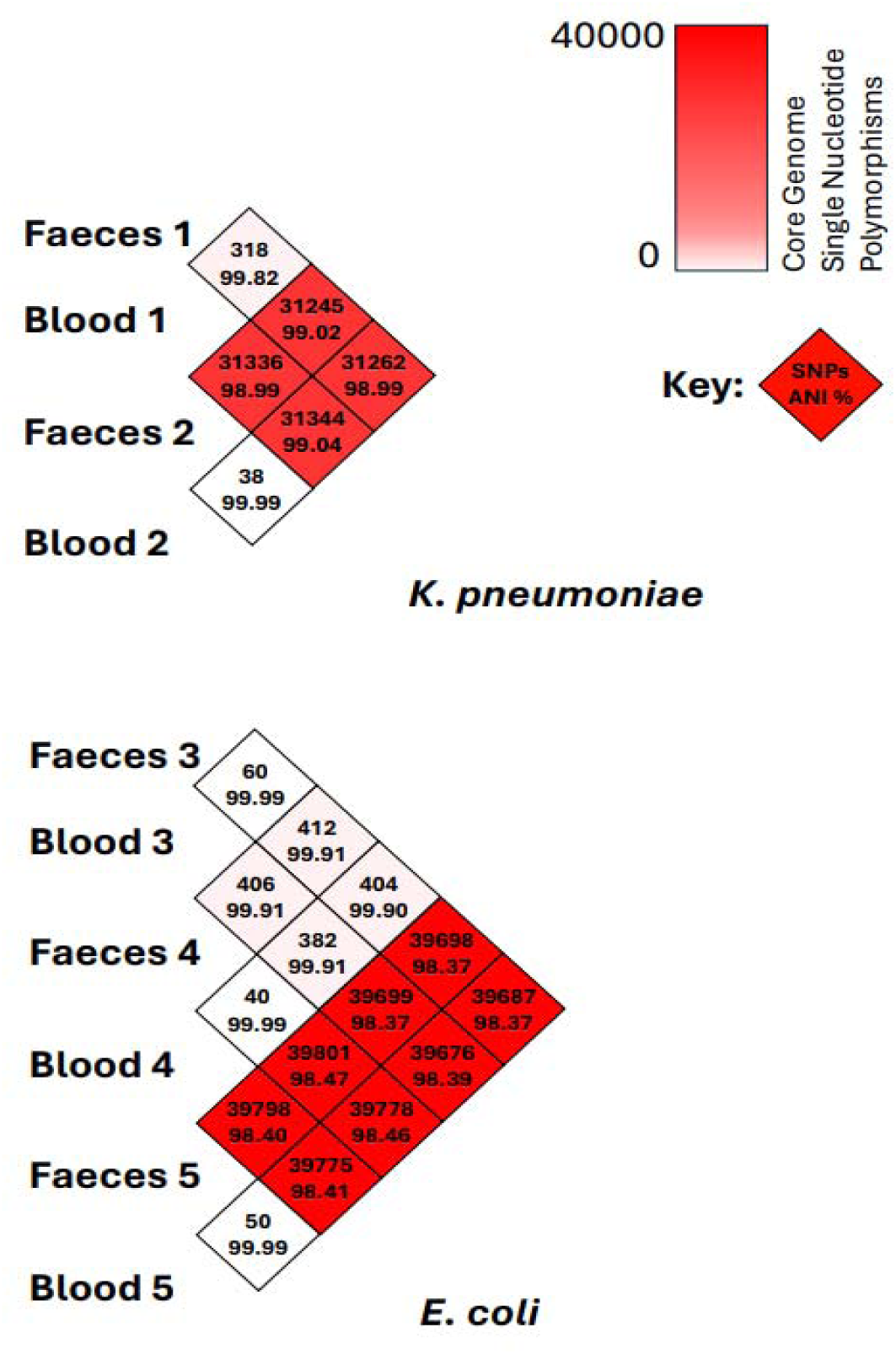
Pairwise comparison of Average Nucleotide Identity and core genome SNPs between isolates

Isolates were next compared for core genome SNP differences (Figure 1B.) For *K. pneumoniae* isolates, Pair 2 exhibited the lowest SNP difference (38 SNPs), while Pair 1 showed the highest (318 SNPs). In *E. coli* isolates, within-pair SNP distances ranged from 40 SNPs in Pair 4 to 60 SNPs in Pair 3. These findings were in alignment with the ANI comparison.

### Initial genomic comparison: ARGs, virulence genes and plasmids

At this point, all pairs were considered to be ‘linked’ and an initial genomic comparison of ARGs, virulence genes and plasmids of the faecal and blood isolates within each pair was performed (Figure 2). There were between 1-13 ARGs detected in each isolate, predicted to confer resistance to various drug-classes (Figure 2A). The presence of at least one ESBL-gene was confirmed in each isolate. ARG profiles were identical within pairs, except for Pair 4, in which *bla*TEM-1B was detected in the blood isolate but not the faecal counterpart, suggesting acquisition between the dates of sampling, possibly during GIT-blood translocation.

**Figure 2A:**
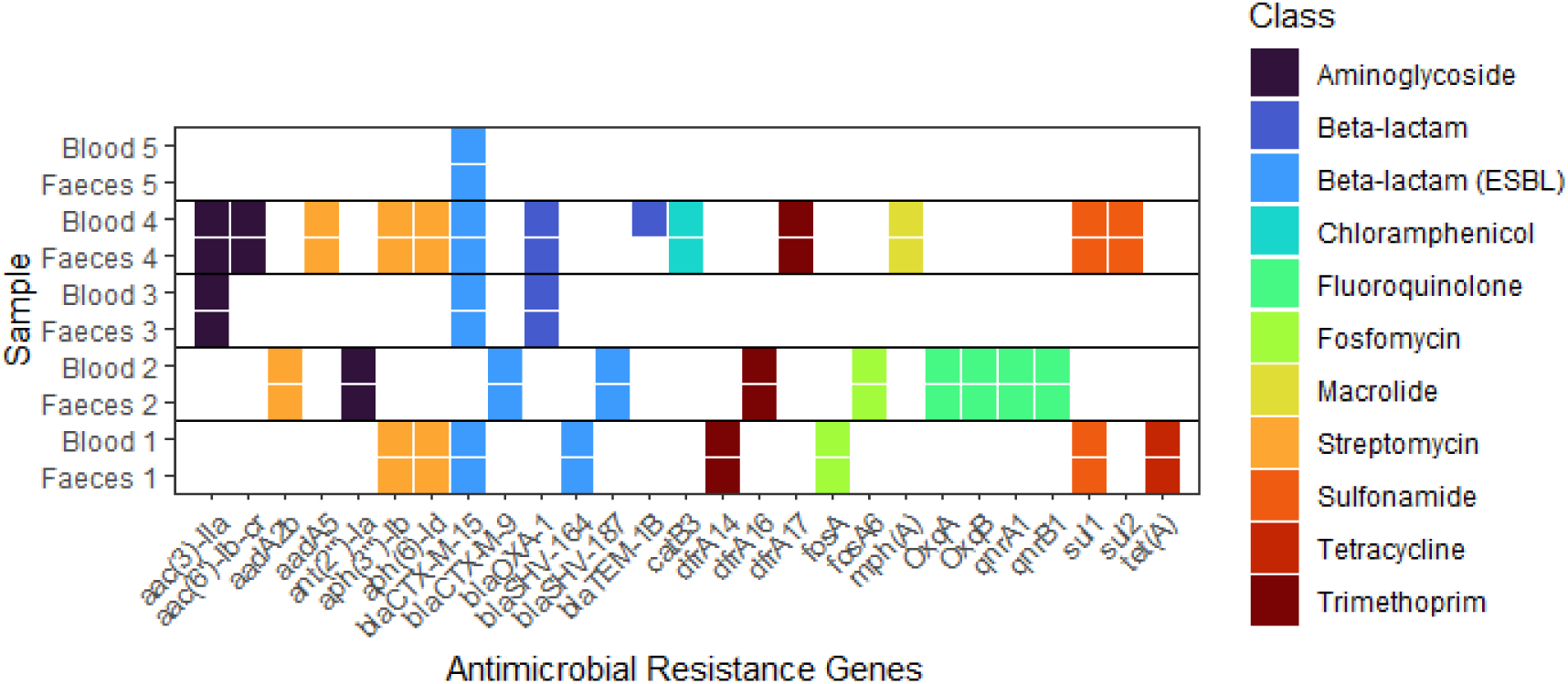
Antimicrobial resistance gene (ARG) presence in all isolates. Each tile represents the presence of an ARG as detected by ResFinder 4.2.2. using whole genome sequencing data. ARGs are categorised by drug class that they are predicted to confer resistance to.

There were between 51-86 virulence genes detected in each isolate, encoding multiple classes of virulence factors (Figure 2B/C). Virulence gene presence was identical between faecal and blood isolates, indicating no loss or acquisition of virulence genes in GIT-blood transition in any pair.

**Figure 2B:**
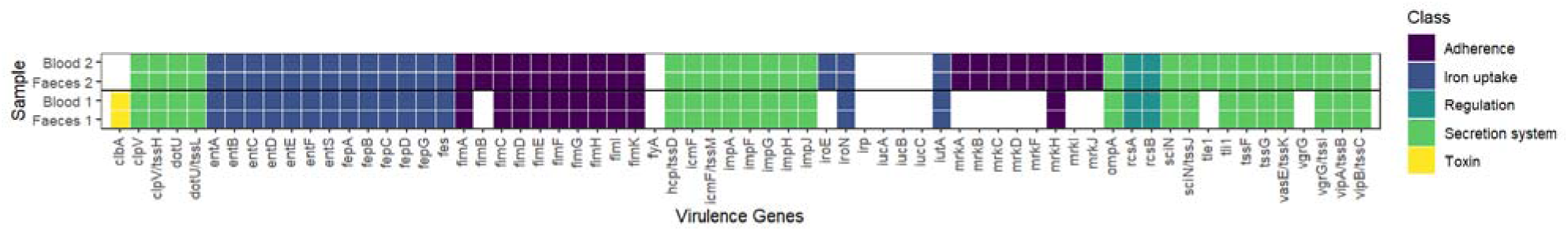
Virulence gene presence in *K. pneumoniae* isolates. Each tile demonstrates the presence of a virulence gene as detected by the VF analyser using whole genome sequencing data. Virulence genes are categorised by class of virulence factor encoded by the gene.

**Figure 2C:**
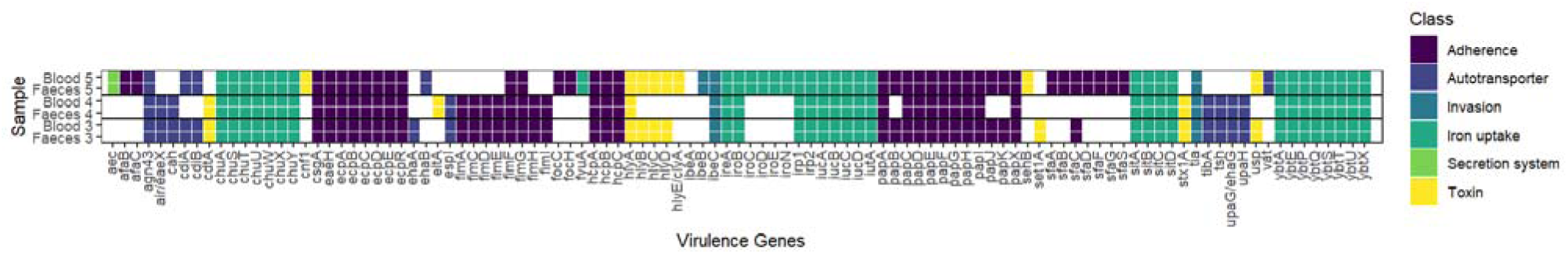
Virulence gene presence in *E. coli* isolates. Each tile demonstrates the presence of a virulence gene as detected by the VF analyser using whole genome sequencing data. Virulence genes are categorised by class of virulence factor encoded by the gene.

There were between 3-8 plasmids detected in each isolate (Figure 2D). Whilst the majority of pairs demonstrated identical plasmid profiles, indicating stable plasmid carriage during the likely GIT-blood transition, Pair 1 demonstrated loss of IncFIB(K) from the faecal isolate and gain of IncN and IncFII(Yp) plasmids in the blood isolate.

**Figure 2D:**
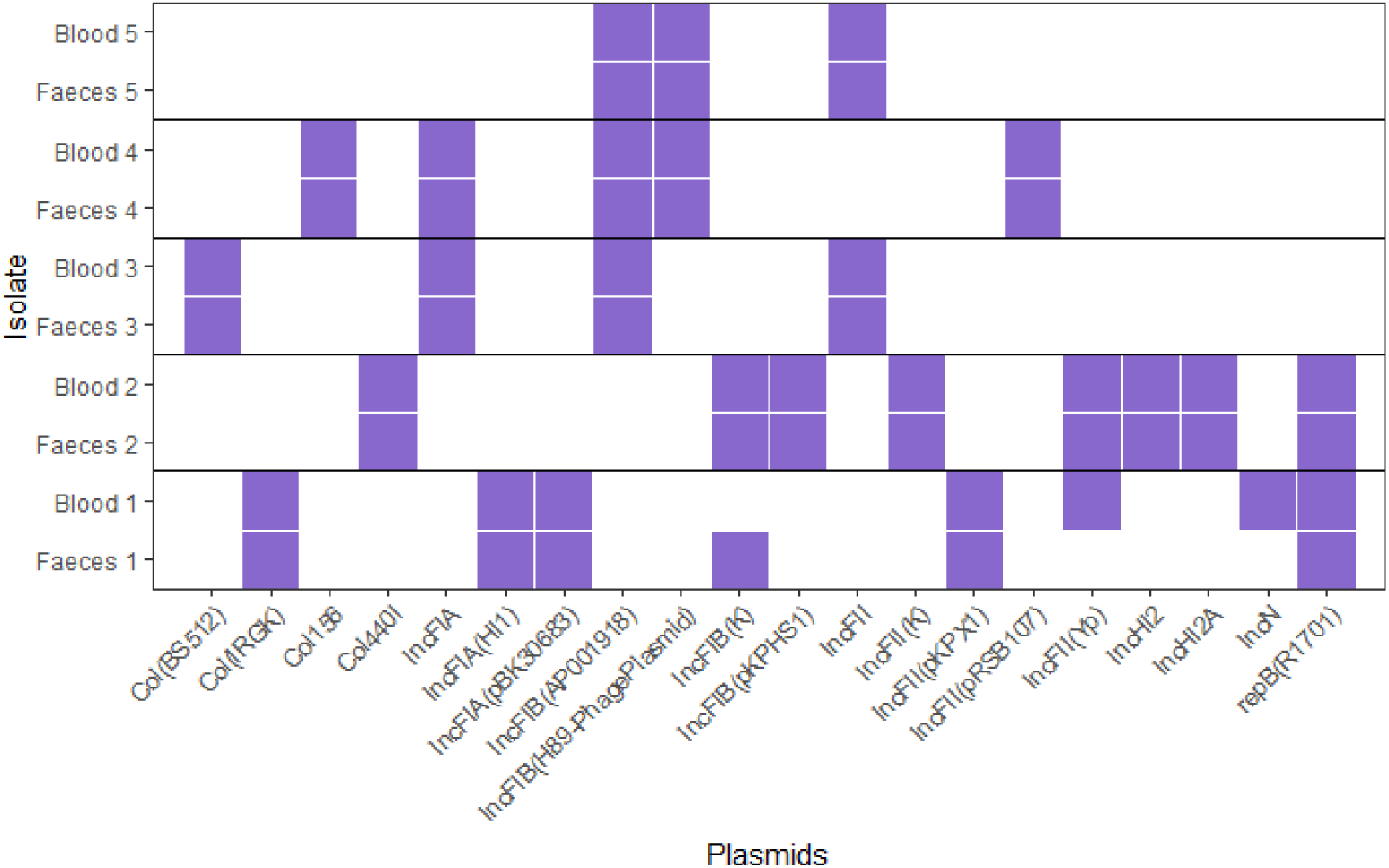
Plasmid presence in all pairs of isolates. Each tile demonstrates the presence of a Plasmid replicon gene in each isolate as detected by PlasmidFinder, using whole genome sequencing data.

### Phenotypic comparison: Antimicrobial resistance

Disc diffusion testing was used to screen for a change in antimicrobial resistance between faecal and blood isolates of each pair. A change in resistance phenotype was identified for three drug-bug combinations in 2/5 pairs (Supplementary figure 1).

There was also a statistically significant difference in zone size between faecal and blood isolates that did not cross the EUCAST clinical breakpoint for four drug-isolate combinations in 2/5 pairs. (Supplementary figure 2, Supplementary table 1).

These within-pair differences were only confirmed by Broth microdilution in two pairs. Pair 2 demonstrated an increase in MIC for Ciprofloxacin, Piperacillin-Tazobactam, Cefotaxime and Gentamicin in the blood isolate compared to its faecal counterpart, indicating increased resistance development to these four antimicrobials between the two sampling timepoints. (Figure 3A). In Pair 3, MIC was decreased for Piperacillin-Tazobactam in the blood isolate compared to faecal, indicating increased susceptibility development in GIT-blood transition. (Figure 3B).

**Figure 3A:**
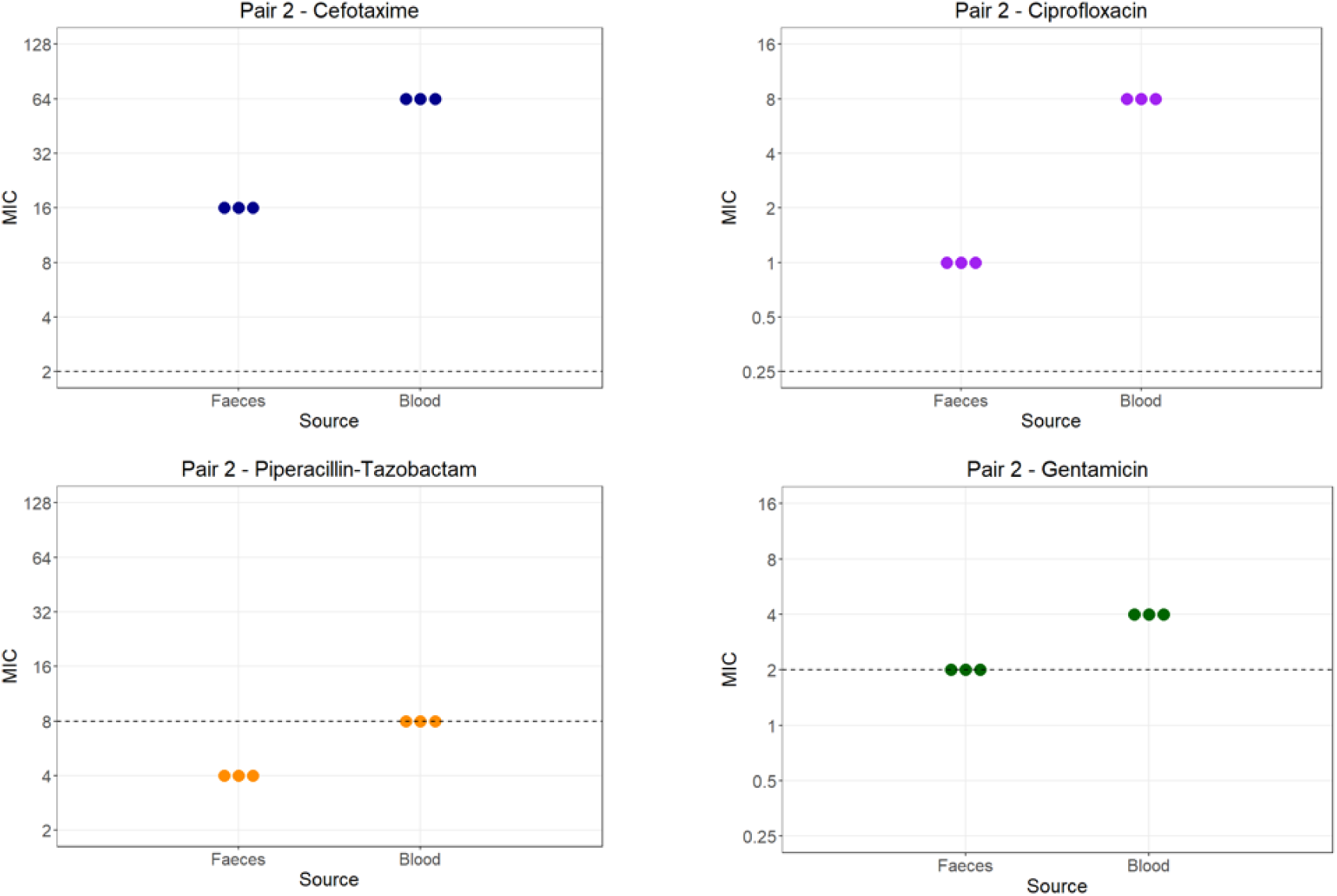
Minimum inhibitory concentration (MIC) (µg/ml) as determined by broth microdilution of Ciprofloxacin, Co-amoxiclav, Piperacillin-Tazobactam, Cefotaxime and Gentamicin for Pair 2. Each plotted point represents the MIC (µg/mL) of a single biological replicate in broth microdilution. Dashed line = EUCAST clinical breakpoint for MIC.

**Figure 3B:**
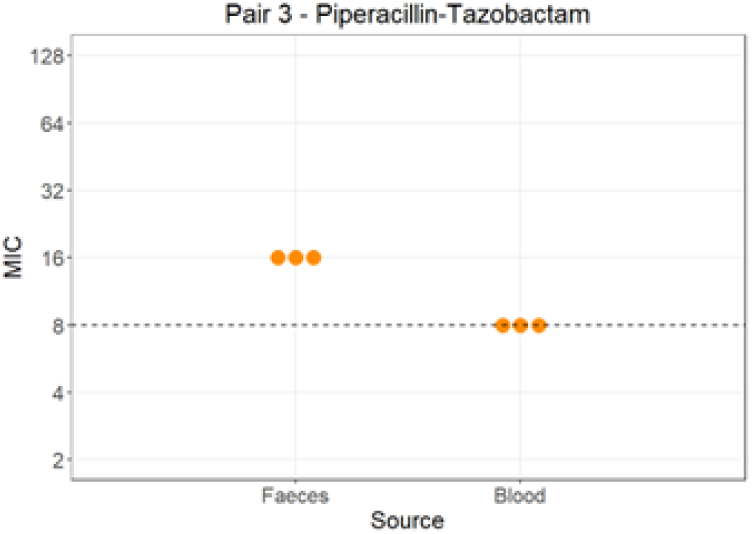
Minimum inhibitory concentration (MIC) as determined by broth microdilution of Piperacillin-Tazobactam for Pair 3. Each plotted point represents the MIC of a single biological replicate in broth microdilution. Dashed line = EUCAST clinical breakpoint for MIC.

### Phenotypic comparison: Biofilm formation

Because all patients were found to be diagnosed with CLABSI, and the importance of biofilm formation in CLABSI pathogenesis, we investigated for any difference in biofilm formation between faecal and blood isolates in each pair. In 4/5 pairs, blood isolates demonstrated significantly increased biofilm formation in comparison to the faecal counterpart, as determined by a paired t-test. (Figure 4). All p-values are stated in Supplementary table 2. Overall, the isolates in Pair 2 demonstrated the strongest biofilm formation.

**Figure 4:**
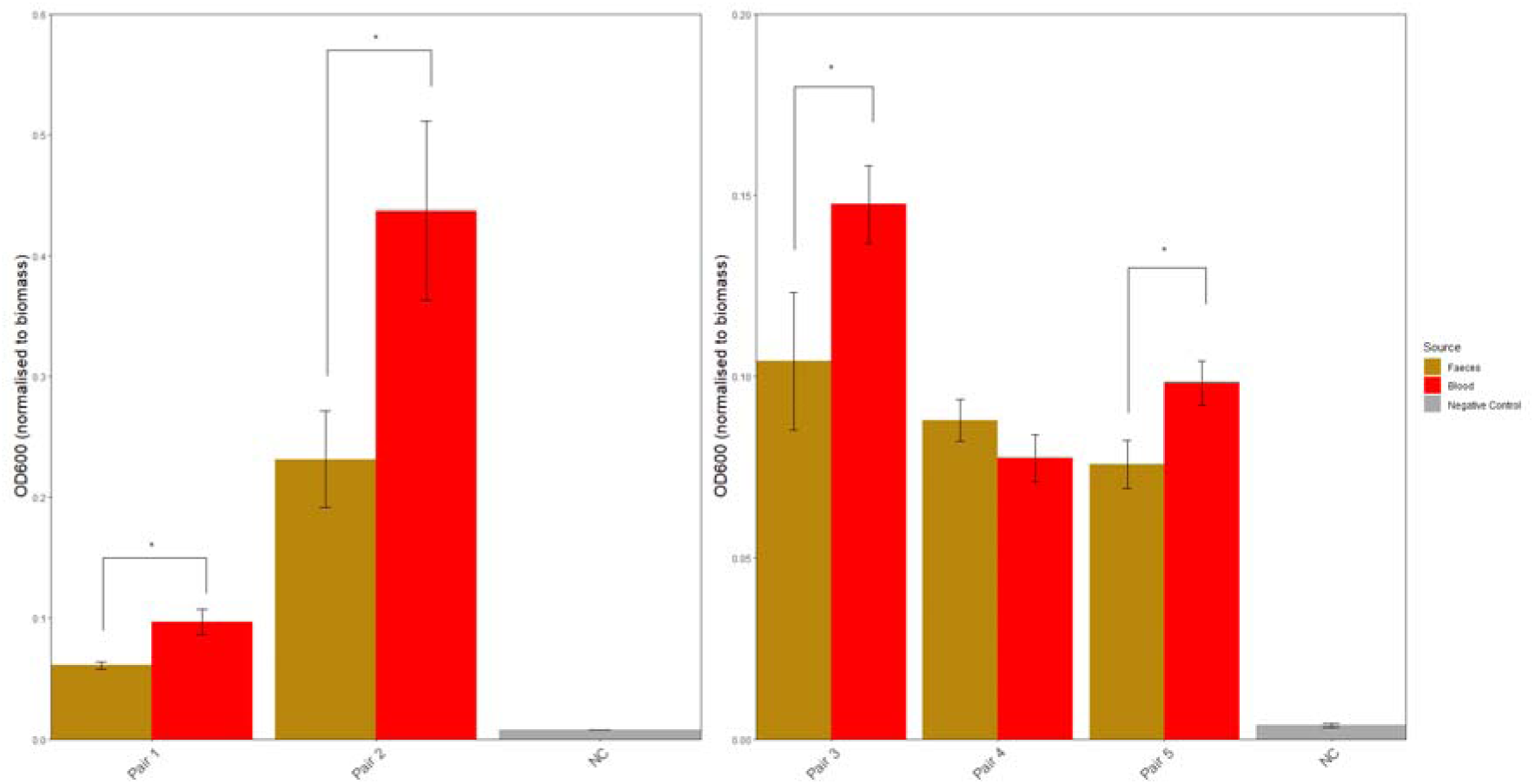
Crystal violet assay for biofilm formation in 96-well plate. Each bar represents mean OD_600_ (n=12) for faecal and blood isolates after crystal violet staining, normalised to biomass. This is used as the measure of biofilm formation capacity. Error bars represent standard error of the mean. * = Statistical significance as determined by paired t-test, p=0.05.

### Comparative genomics: Breseq analysis

Breseq was used to identify mutations in each pair. Most mutations were Indels, and the majority of these were ‘Function unknown’ or ‘Unclassified.’ (Supplementary figure 3) Mutations occurring within 100bp of genes with a known function were assigned to COG categories (Figure 5). There were six categories in which mutations occurred in all pairs. Of these, most occurred in ‘L: Replication, recombination and repair,’ ‘E: Amino acid transport and metabolism,’ and ‘G: Carbohydrate transport and metabolism.’

**Figure 5:**
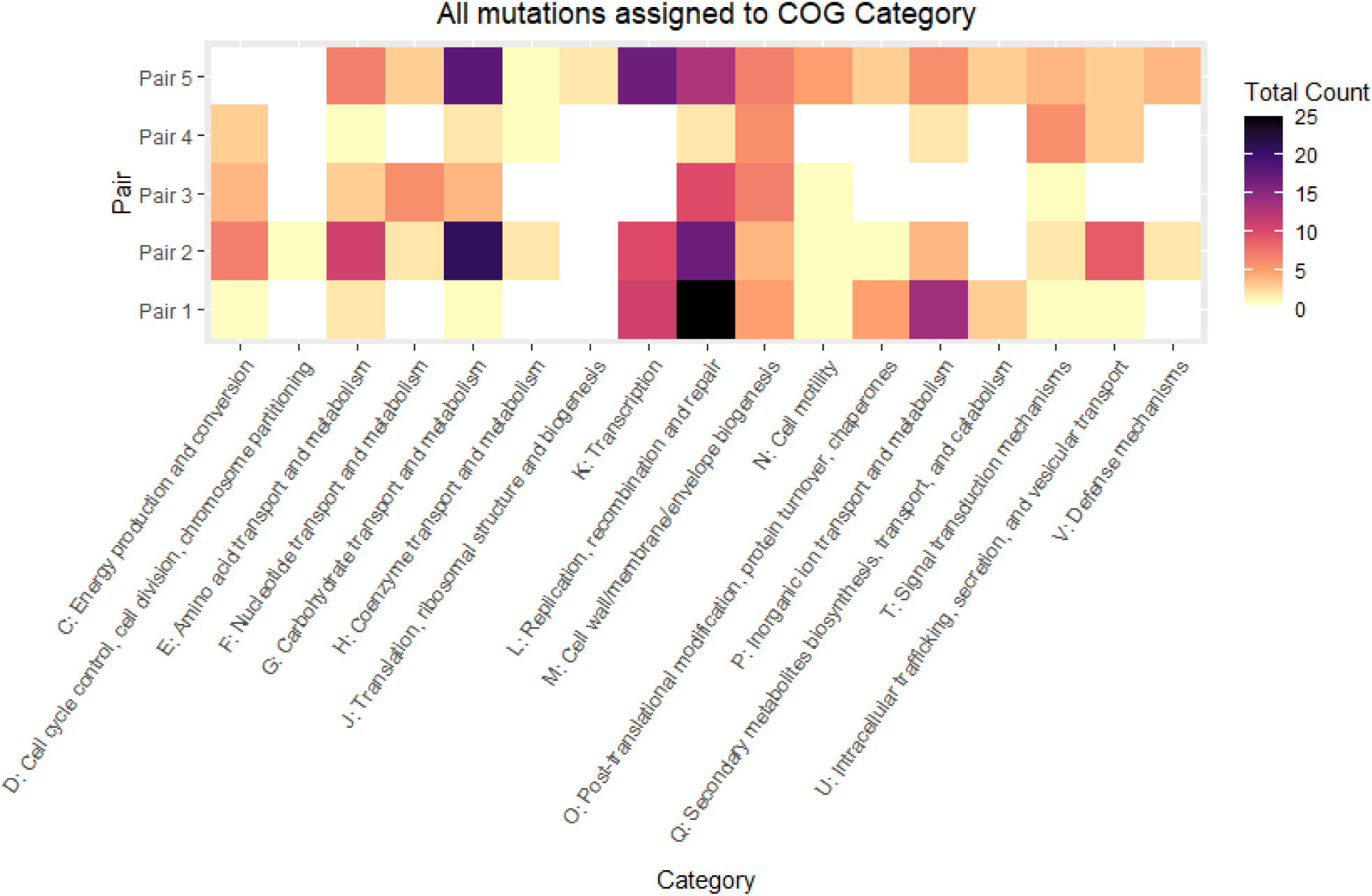
Heatmap of all mutations assigned to COG categories across isolate pairs. Each tile represents the total count of mutations within each COG (Cluster of Orthologous Groups) category for each isolate pair. White tiles represent the absence of mutations in a specific COG category.

No mutations were identified in specific ARGs identified by ResFinder, however four pairs demonstrated mutations in genes encoding efflux pumps and transporter proteins involved in AMR (Table 3). Four pairs demonstrated mutations in at least one known virulence factor gene. Two pairs demonstrated mutations in or related to previously reported bloodstream survival factors. All pairs demonstrated a variety of mutations in genes related to biofilm formation. In Pair 1, there was evidence of acquisition of another copy of the *hha* gene, along with modification of the Shine-Delgarno intergenic region (Figure 6)

**Figure 6:**
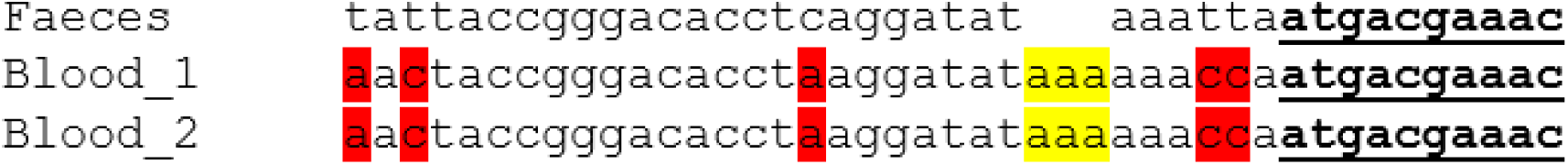
Alignment of upstream intergenic region and first 10bp (underlined) of *hha* gene of faecal isolate and two copies in blood isolate. (Yellow = duplication of bp. Red = intergenic SNP).

**Table 3:** Specific mutations identified in relation to phenotypes of interest.

| Pair | Mutation type | Mutation | Annotated gene | Relevance of mutation |
| --- | --- | --- | --- | --- |
| 1 | nsSNP | D47E (GAC→GAA) | ompK36 | Bloodstream survival factor |
|  | nsSNP | Q62K (CAA→AAA) | dam | Bloodstream survival factor |
|  | nsSNP | T57A (ACT→GCT) | dam | Bloodstream survival factor |
|  | nsSNP | Q141L (CAG→CTG) | dam | Bloodstream survival factor |
|  | Insertion | +GGT | Haemolysin expression modulating protein/hypothetical protein | Regulator of virulence factors, known to regulate biofilm formation |
|  | iSNP | 2 bp→TT | Haemolysin expression modulating protein/hypothetical protein | Regulator of virulence factors, known to regulate biofilm formation |
|  | iSNP | G→T | Haemolysin expression modulating protein/hypothetical protein | Regulator of virulence factors, known to regulate biofilm formation |
|  | iSNP | A→G | Haemolysin expression modulating protein/hypothetical protein | Regulator of virulence factors, known to regulate biofilm formation |
|  | iSNP | A→T | Haemolysin expression modulating protein/hypothetical protein | Regulator of virulence factors, known to regulate biofilm formation |
|  | sSNP | A40A (GCT→GCC) | Haemolysin expression modulating protein | Regulator of virulence factors, known to regulate biofilm formation |
|  | sSNP | R11R (CGC→CGG) | Haemolysin expression modulating protein | Regulator of virulence factors, known to regulate biofilm formation |
|  | Deletion | Δ916 bp | Fec operon | Iron uptake and utilisation |
| 2 | nsSNP | T37I (ACT→ATT) | Multiple antibiotic resistance protein MarB | Drug-efflux pump implicated in antibiotic resistance |
|  | Deletion | Δ348 bp | YcAD - MFS transporter | Drug efflux pump implicated in antibiotic resistance |
|  | Deletion | Δ254 bp | permease of drug/metabolite transporter | Drug efflux pump implicated in antibiotic resistance |
|  | Deletion | Δ250 bp | inner membrane protein/drug antiporter | Drug efflux pump implicated in antibiotic resistance |
|  | Deletion | Δ243 bp | LysR | Transcriptional regulator implicated in antibiotic resistance |
|  | Deletion | Δ241 bp | MFS-type transporter | Drug efflux pump implicated in antibiotic resistance |
|  | Deletion | Δ239 bp | RND efflux transporter | Drug efflux pump implicated in antibiotic resistance |
|  | Deletion | Δ236 bp | LysR transcriptional regulator | Drug efflux pump implicated in antibiotic resistance |
|  | Deletion | Δ321 bp | acrR | Transcriptional regulator implicated in antibiotic resistance and biofilm formation |
|  | nsSNP | A434S (GCC→TCC) | Protease III precursor (EC 3.4.24.55) | Biofilm formation |
|  | nsSNP | V701A (GTG→GCG) | T6SS component TssM (IcmF/VasK) | Virulence factor, involved in biofilm formation |
| 3 | Deletion | Δ235 bp | Lipopolysaccharide ABC transporter | Transporter protein implicated in antibiotic resistance |
|  | nsSNP | T115A (ACC→GCC) | Phage endopeptidase Rz | Phage protein, adjacent to virulence factor iss (involved in bloodstream survival) |
|  | nsSNP | L237P (CTT→CCT) | Mobile element protein | MGE adjacent to Fec operon (involved in iron uptake and utilisation) |
|  | nsSNP | A238G (GCC→GGC) | Mobile element protein | MGE adjacent to Fec operon (involved in iron uptake and utilisation) |
|  | nsSNP | S932T (AGC→ACC) | Antigen 43 | Virulence factor, involved in biofilm formation |
|  | nsSNP | A763V (GCC→GTC) | Antigen 43 | Virulence factor, involved in biofilm formation |
|  | sSNP | G765G (GGA→GGT) | Antigen 43 | Virulence factor, involved in biofilm formation |
| 4 | Deletion | Δ286 bp | MFS type transporter | Drug efflux pump implicated in antibiotic resistance |
|  | sSNP | G977G (GGG→GGT) | Antigen 43 | Virulence factor, involved in biofilm formation |
|  | sSNP | G972G (GGT→GGA) | Antigen 43 | Virulence factor, involved in biofilm formation |
|  | nsSNP | S932T (AGC→ACC) | Antigen 43 | Virulence factor, involved in biofilm formation |
|  | nsSNP | V375A (GTC→GCC) | Antigen 43 | Virulence factor, involved in biofilm formation |
|  | Deletion | Δ265 bp | Ferrochelatase | Iron uptake and utilisation |
|  | Deletion | Δ236 bp | Ferredoxin reductase | Iron uptake and utilisation |
|  | Deletion | Δ119 bp | cog3210 | Iron uptake and utilisation |
|  | Deletion | Δ95 bp | cog3210 | Iron uptake and utilisation |
| 5 | Synonymous SNP | L123L (CTG→TTG) | Manganese ABC transporter, inner membrane permease protein SitC | Iron uptake and utilisation |
|  | Deletion | Δ297 bp | Transcriptional regulator, acrR family | Transcriptional regulator implicated in antibiotic resistance and biofilm formation |
|  | Deletion | Δ295 bp | Spermidine/putrescine ABC transporter | Transporter protein implicated in antibiotic resistance |
|  | Deletion | Δ281 bp | Fe2+ ABC transporter, substrate binding protein | Iron uptake/homeostasis |
|  | Non-synonymous SNP | V32E (GTG→GAG) | Transcriptional repressor of aga operon | Transcriptional regulator involved in biofilm formation |
|  | Non-synonymous SNP | M300L (ATG→CTG) | Lipopolysaccharide export system permease protein LptG | Biofilm formation |
|  | Deletion | Δ246 bp | TctA transporter | Drug-efflux pump implicated in antibiotic resistance |
|  | Deletion | Δ245 bp | AaeAB efflux system | Drug efflux pump implicated in antibiotic resistance |

## Discussion

In this study, we compared blood isolates with the previously acquired faecal isolates from 5 children with GNBSI, and in each instance demonstrated concordance in sequence type and species-specific typing methods. This is highly suggestive that the faecal-blood isolate pairs were genomically ‘linked’ and that the invasive blood isolate was likely to have been acquired from the GIT. Average nucleotide identity has been used as a marker of genomic relatedness in previous studies comparing faecal-blood isolates in children [22, 23] but there is no defined consensus on a threshold that indicates isolates as similar enough to be linked. Thresholds of 99.99% have been previously used to indicate isolates being the same strain [24] Using this threshold, 4/5 pairs (Pairs 2-5) shared ANI values convincing of being the same strain. The lower ANI in Pair 1 (99.82%) could be somewhat explained by the identified plasmid changes in this pair. This is consistent with observations that MGE movement can lead to a lower ANI in isolates that are highly likely to be concordant, as noted by Vornhagen *et al*. when comparing wzi-type concordant rectal and blood *K. pneumoniae* isolates [25]. However, this finding could also be indicative of the blood isolate having evolved to be a different clade of the same sequence type as the faecal isolate in this pair, as indicated by the higher number of core genome SNPs calculated in further analysis.

The number of core genome SNPs between faecal and blood isolates in each pair ranged from 38 SNPs (Pair 2) to 318 SNPs (Pair 1), indicating varying degrees of evolutionary distance between faecal and blood isolates in each patient. It is possible that our BSI isolates, although related, are not immediate descendants of the cognate faecal isolate and that there could have been closer ancestors in the GIT which were not picked up during sampling. Similarly, there may have been evolution within the blood giving rise to multiple descendants, of which we isolated one. A previous study that identified concordant faecal and blood isolates in patients with *E. coli* bloodstream infections noted that the number of core genome SNPs between faecal and blood isolates was 0 [26]. Another study comparing faecal and blood *K. pneumoniae* isolates found a core genome SNP difference of 6 SNPs between isolates considered to be concordant strains in one patient and 76 SNPs between concordant isolates from another patient. [27]. In our study, the number of SNPs we found is generally higher, and was not proportional to the number of days between faecal and blood isolation. This may represent varied and convoluted routes taken from the GIT to the blood e.g. via other body tissues in each case of GNBSI, requiring adaptation to multiple niches – as opposed to direct GIT-blood translocation Alternatively, the higher number of core genome SNPs in Pair 1 could be considered to contradict the hypothesis that this represents a ‘linked’ pair. However, as with ANI, there is no SNP threshold that is universally agreed as defining isolates as ‘linked’. For example, whilst one study used a threshold of 17 core genome SNPs for this, another study looking at persistence of *E. coli* isolates between GI and urinary tracts used a threshold of ≤500 core genome SNPs [26, 28]. These differences could also be due to differences in sequencing and bioinformatic technologies, as well as the reference genomes used in different studies. Such differences in methodology are recognised as a key barrier to the determination of a defined threshold for strains similarity [29–31]. In addition, the expected rate of mutation of isolates moving between different body niches is currently unknown and unpredictable [32], adding to this challenge. When comparing isolates from the two different patients with *K. pneumoniae* infection, there are over 30,000 core genome SNPS between each. A similar number is seen when comparing the *E. coli* isolates of Patient 5 to both Patient 3 and Patient 4. However, the number of core genome SNPs between Patient 3 and 4 are more comparable to those seen within the pair of Patient 1. This raises suspicion for a common source of colonisation of these patients within the intensive care unit during the time period the isolates were taken. However granular data of overlapping hospital admission times is unavailable.

ARG profile and subsequently AMR phenotypes were compared within ‘linked’ pairs. The transition from GIT colonisation to blood invasion was associated with acquisition of an ARG in 1/5 pairs, (Pair 4) - and in this case, the acquisition of *bla*TEM-1B did not lead to increased resistance to any of the beta-lactam/inhibitor combinations tested. Only in one pair, (Pair 2) did we observe that increases in resistance to any antimicrobials occurred between the two sample timepoints. This is despite that in all five cases, the patient from whom isolates were obtained received multiple antimicrobials during hospital admission. In contrast, for Pair 3, increased susceptibility to Piperacillin-tazobactam was noted. These findings may reflect varying selection pressure exerted by dose and duration of antimicrobial therapy, in balance with environmental adaptation. Unfortunately, the study is limited by lack of granular data on antimicrobial exposure.

These phenotypic changes in AMR are not explained by acquisition or loss of ARGs (Figure 2A). However, by using Breseq for whole genome comparison, we identified deletion of drug-efflux pump repressors that could putatively explain this observation. Several drug-efflux pumps are conserved across Gram-negative species and implicated in AMR. These include MFS-pumps, RND-pumps, and ABC-transporters. The *acrAB*-*tolC* efflux pump is an RND-pump implicated in resistance to multiple antimicrobials. [33]. It has multiple regulatory pathways, including the *acrRAB* operon, in which *acrR* is a repressor, and the *marAB* operon, with *marB* as a repressor. In Pair 2, *marB* and *acrR* deletions were identified (Table 3), which could lead to increased efflux pump activity, explaining the changes in susceptibility. The deletion of *acrR* is particularly interesting because a previous study reported that *acrR* knockout in *K. pneumoniae* isolates resulted in increased MIC to the same antimicrobial classes as seen here [34]. Previous literature also reports an association between *acrR* disruption with a MDR phenotype [35, 36]. However, it must be noted that the *acrR* deletion was identified in Pair 5 but without a change in susceptibility, possibly explained by the variation in other efflux pump changes seen in each pair.

In 5/5 cases in this study, GIT-blood transition was not associated with the acquisition of virulence genes when comparing blood isolates to the presumed faecal ancestor. Because all cases of GNBSI in this study were cases of CLABSI, biofilm formation was chosen as a virulence phenotype of interest. The finding that 4/5 blood isolates exhibited an increase in biofilm formation compared to faecal counterparts, suggests in-host adaptation towards increased biofilm formation may be implicated in the pathogenesis. As noted, the observed phenotypic changes are not explained by acquisition of virulence genes involved in biofilm formation, but comparative genomic analysis using Breseq again identified a variety of putative mechanisms that could potentially underpin our phenotypic observation.

In Pair 2, the *acrR* deletion explored above could also be responsible for increased biofilm formation, as *acrAB*-*tolC* has a central role in biofilm formation in *E. coli* and *K. pneumoniae.* [37, 38]. Studies also link biofilm formation, AMR and *acrAB*-*tolC* activity. In a comparison of *K. pneumoniae* urinary isolates, Vuotto *et al*. observed that those with the highest biofilm forming-capacity also demonstrated the most AMR, correlating with increased *acrAB* activity [39]. Although only a sample of three isolates, this strikingly mirrors the findings of the *K. pneumoniae* blood isolate of Pair 2, raising the possibility of a single intra-host adaptation contributing to both antimicrobial resistance and biofilm formation. Hennequin *et al* demonstrated this *in vitro*, reporting that sub-MIC cephalosporin exposure in *E. coli* isolates resulted in enhanced *acrAB* expression and biofilm formation [40]. This could be an example of antimicrobial exposure selecting for a resistance-conferring mutation that also increases virulence and propensity to invasive infection. If seen consistently in a larger cohort, this could suggest a promising interventional target. For example, RND-efflux pump inhibitors have been explored for use as antimicrobial-adjuvant therapy for resistant infections, and for biofilm disruption in invasive infection [41].

Furthermore, in this pair, nsSNPs in two biofilm-related virulence genes, *IcmF* and Protease III precursor were identified, possibly contributing to the phenotypic change. Similarly, in Pair 3, two nsSNPs in the biofilm-associated virulence gene ‘antigen 43’ were noted. Mutations in this gene were noted when Nielson *et al* compared ‘linked’ urine and faecal isolates, and hypothesised this related to increased biofilm formation [42].

Pair 1 exhibited a potential link between biofilm formation and plasmid acquisition. This pair underwent loss of IncFIB(K) and acquisition of IncFII(Yp) between faecal and blood isolates (Figure 2D), Breseq analysis revealed a duplication of the *hha* gene in the blood isolate compared to the faecal isolate (Table 3), with the additional copy residing on a contig containing multiple IncF-related genes, suggesting plasmid-mediated acquisition. The *hha* gene encodes haemolysin-expression-modulating protein and is a regulator of virulence factors involved in cell adhesion and biofilm formation in Gram-negative species. Studies in *E. coli* have identified *hha* as a positive regulator of biofilm formation and of virulence [43, 44]. This is less studied in *K. pneumoniae*, however Bandeira *et al* described *hha* presence to be associated with *K. pneumoniae* biofilm-forming isolates, particularly on indwelling central lines [45]. Plasmid-mediated acquisition of *hha* was described by Krall *et al* who compared ‘linked’ faecal and blood isolates finding that *hha* acquisition on an IncFII plasmid increased the invasiveness of *E. coli* isolates in a gut-organoid model [46]. They concluded *hha* was involved in GIT-blood adaptation but did not explore biofilm formation specifically. Here, one could similarly hypothesise that IncFII plasmid-mediated acquisition of *hha* could be an adaptive change leading to increased biofilm formation and procession to GNBSI.

Other studies report *hha* has a negative influence on biofilm formation [47, 48] which this hypothesis contradicts. Interestingly, in both copies of the *hha* gene in the blood isolate, a 3bp GGT insertion was noted 1bp downstream of *hha*. Upon gene alignment, this appeared to be an AAA duplication, disrupting the Shine-Dalgarno sequence 7bp before the gene (Figure 6). This could potentially reduce the efficiency of protein translation. Hence, one could equally hypothesize, assuming a negative-regulatory role of *hha*, that after plasmid acquisition during GIT colonisation, the compensatory mutation in the Shine-Dalgarno region led to increased biofilm formation and aided bloodstream invasion. Further investigation is required to clarify the role of *hha* in invasion and the combined effect of the observed SNPs, Shine-Dalgarno mutation, and *hha* copy number on biofilm formation.

The transition from GIT-blood, and subsequent survival in the bloodstream will involve the ability to adapt to a harsh and dynamic environment, with factors such as varying nutrient abundance and host immunity to contend with. As noted, this adaptation did not appear to result from acquisition of virulence genes in these five cases, but was associated with varied changes across the genome. No single specific mutation occurred in all pairs, likely reflecting the differing clinical scenarios and selection pressures driving diverse evolutionary trajectories en-route to bloodstream infection. However, in all pairs, most mutations occurred in the COG category ‘Replication, recombination and repair’, including a mutation in *dam* a known bloodstream survival factor, in Pair 1 [49]. These genes are involved in environmental and stress response and selection for mutations could be expected in adaptation to challenging environments, for example maintaining efficient DNA repair mechanisms in the presence of reactive oxygen species, host immune defences or nutrient variability. Another commonality was mutations in ‘cell wall/membrane/envelope biogenesis,’ ‘cell motility,’ and ‘signal transduction’ categories. Such mutations might confer altered chemotaxis or adherence, allowing tissue invasion. Few mutations were observed in ‘Cell cycle control and repair’ and ‘Translation, ribosomal structure and biogenesis.’ This demonstrates the strong selective pressure to conserve key cellular processes integral to bacterial survival during the transition between GIT and blood environments. Many mutations were classified as ‘Function Unknown’ within COG categories but may hold significance. For example, in Pair 3, a non-synonymous SNP was identified in a phage endopeptidase gene adjacent to the gene *iss.* The *iss* gene has is associated with bloodstream survival in *E. coli* through mediating serum and complement resistance, and increased mRNA copy number has been associated with increased serum survival [50, 51]. Mutations in the adjacent phage endopeptidase could influence the activation or expression of *iss*, with effects on bloodstream survival. This represents a potential avenue for further research.

When comparing blood isolates to their faecal ancestors using Breseq, we noted mutations related to carbohydrate and amino acid transport in 5/5 pairs. This included an nsSNP in *ompk36* in Pair 1, a transport protein also previously identified as a bloodstream survival factor in *Klebsiella pneumoniae* [52]. Further, we identified mutations related to iron uptake utilisation in 4/5 pairs. Iron availability and chemical form varies in different tissues and a ‘functional hierarchy’ whereby iron uptake systems are utilised variably across environments has been discussed by Garcia *et al* in the context of UTI [53]. The findings of *fec* operon deletion in Pair 1 and the transposase mutation near the *fec* operon in Pair 3 may suggest a genomic shift away from ferric citrate uptake, whilst the *sitC* virulence gene mutation and ferrous (Fe^2+^) ABC transporter deletion in Pair 5 may suggest a shift away from Fe^2+^ uptake. These mutations likely reflect adaptation towards iron-acquisition in the bloodstream, where ferric citrate is less abundant than the gut and Fe^2+^ is less abundant due to oxygenated conditions. Further exploration could inform therapeutic strategies. For example, *tonB* which regulates the *fec* operon has been explored as a potential drug target against *E. coli* [54]. However, this may not be an appropriate target in the context of bloodstream infection if functional redundancy of the *fec* operon is a common adaptation.

The strengths of this study include the combination of genomic with phenotypic comparison, using isolates from the same patient to gain insight into within-host evolution and link phenotypic differences to changes at the genomic level. To our knowledge, there are few studies in the existing literature that have investigated this. Further, we compared mutations across the whole genome, rather than focusing only on known ARGs and virulence genes identifying potentially relevant mutations that would have been missed by a narrower approach. Limitations of our study lie in the fact that it cannot be definitively concluded that these pairs are ‘linked’ given the lack of an existing robustly defined threshold for strain similarity/genomic relatedness - this in itself highlighting an area for further work. In addition, the retrospective nature and ability to only include five cases of GNBSI, without granular clinical data, meaning the significance and generalisability of these findings cannot be assumed. The use of short-read rather than long-read sequencing makes it harder to discern plasmid-mediated changes that could contribute to adaptation. Further, phenotypic assays for biofilm formation demonstrated changes *in vitro*, but likely do not reflect true clinical conditions.

Whilst this study shows individual instances of mutations of potential clinical relevance in each pair which may represent avenues for further study; it is clear that there is a diverse range of possible evolutionary trajectories in GIT-blood transition, even in this small study, so a common pathway for clinical intervention may be difficult to elucidate. Future study with a larger sample size that used faecal isolates from proven ‘linked’ pairs to colonising isolates that did not achieve bloodstream invasion, could be a more promising avenue to identify pathogen factors e.g virulence gene SNPs that may predispose to bloodstream infection. Previous case-control studies with this approach have been used, but not including WGS data [55]. Analysis of isolates from other body-site isolates would be beneficial, along with information around antimicrobial exposure, clinical severity, and GNBSI source, giving a more comprehensive insight into the relevance of adaptations to differing clinical scenarios, and how findings could translate to clinical intervention.

Overall, this study used genomic and phenotypic comparison of ‘linked’ pairs of blood and faecal *K. pneumoniae* and *E. coli* isolates to investigate evolutionary and adaptive changes relevant to the transition from GIT colonisation to bloodstream invasion and GNBSI. Phenotypic changes in AMR were variable, however biofilm formation and metabolic flexibility were common phenotypic adaptations towards bloodstream infection in this small sample. These changes were not found to be associated with acquisition of ARGs or virulence genes, but rather, with a diverse array of genomic changes, including mutation of virulence genes, efflux pumps and acquisition of plasmid-mediated genes. No single mutation occurred in all pairs, however changes in functional categories of genes involved in transport and metabolism were common. These findings demonstrate a snapshot of the vast evolutionary landscape of intra-host adaptation that is traversed in GNBSI pathogenesis. However, significant further study is needed to assess whether these findings could feasibly translate into clinical intervention for GNBSI treatment or prevention.

## Acknowledgements

APR acknowledges funding from the Medical Research Council, Biotechnology and Biological Sciences Research Council and Natural Environmental Research Council which are all Councils of UK Research and Innovation (grant no. MR/W030578/1) under the umbrella of the JPIAMR (Joint Programming Initiative on Antimicrobial Resistance), and UKRI through the Strength in Places Fund (grant no. SIPF 36348). EA and RP are by the MRC via the LSTM and Lancaster University PhD Doctoral Training Program (Grant no. MR/N013514/1)

