## Supplemental tables and figures for "From Colonisation to Invasion: Genomic and Phenotypic Comparison of Faecal and Bloodstream Isolates from the same patients" (from another version of this preprint: version 1, 2025-07-15)

**Supplementary figures and tables :**

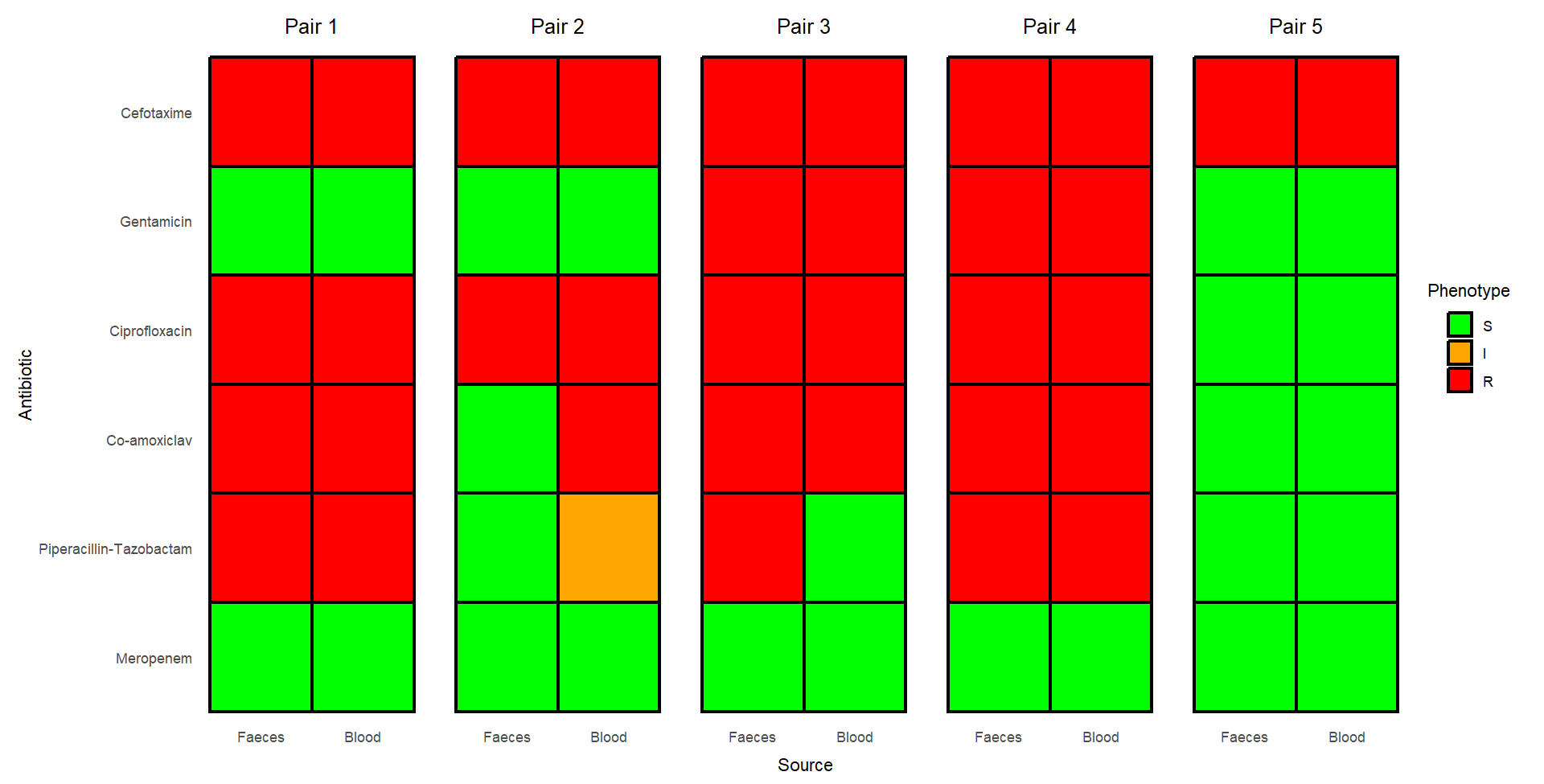

**Supplementary figure 1:** Mode antimicrobial susceptibility phenotype using disc diffusion testing (n=9) against six antimicrobials, as per EUCAST clinical breakpoints. S=Sensitive, I=Sensitive with increased dosing, R=Resistant.

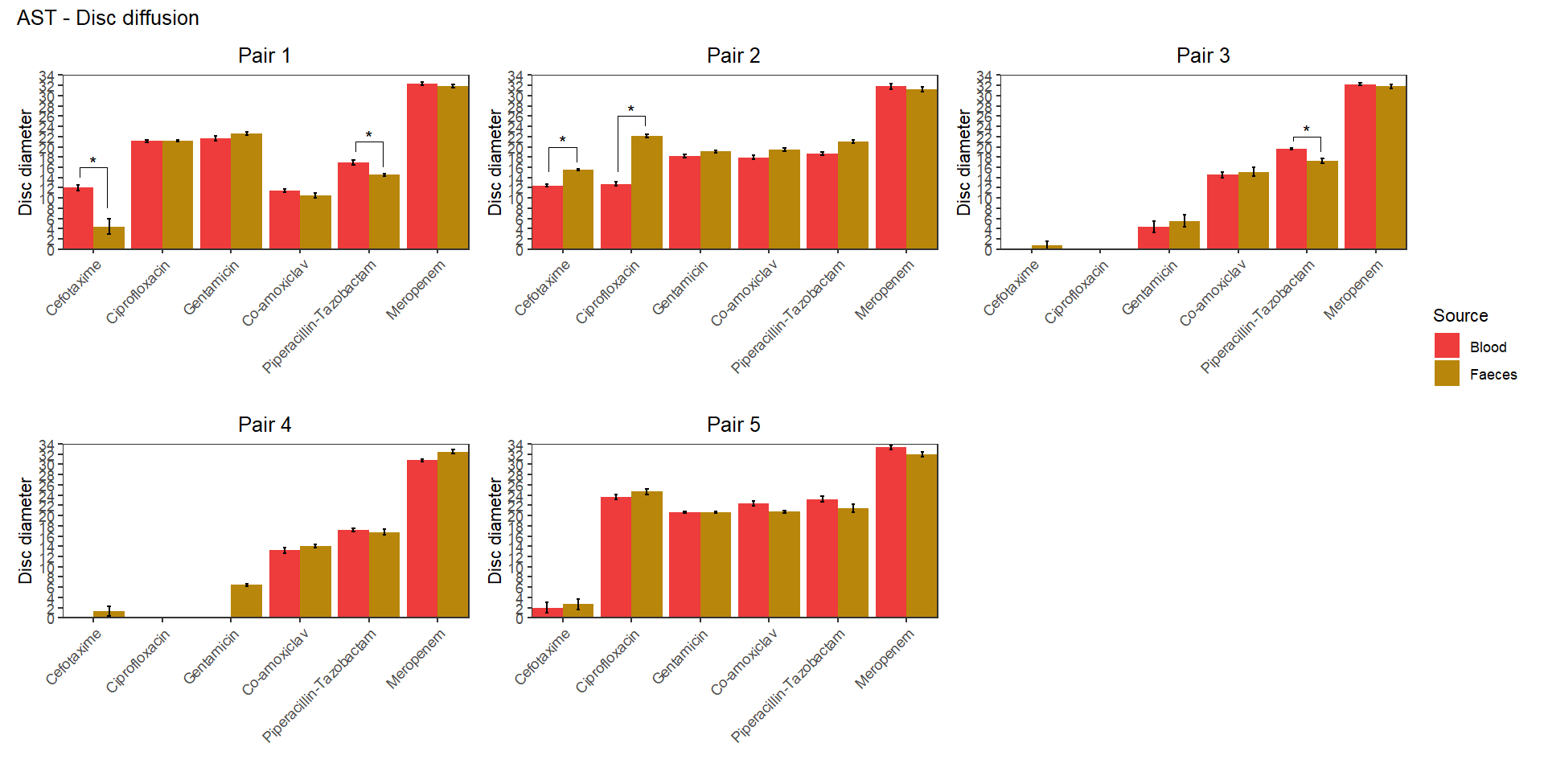

**Supplementary figure 2:** Mean disc diameter (n=9) of faecal and blood isolates tested against six antimicrobials for all 5 pairs. On each chart, each bar represents the disc diameter of faecal and blood isolates against each antimicrobial. Error bars represent standard error of the mean. * = statistical significance as determined by Mann-Whitney U test, p=0.05.

**Supplementary table 1:** Mann-Whitney U Test to ascertain significant difference between faecal and blood isolate disc diameters.

|  | **Pair 1** | **Pair 2** | **Pair 3** | **Pair 4** | **Pair 5** |
| --- | --- | --- | --- | --- | --- |
| **Antibiotic** | **P value** | | | | |
| **Cefotaxime** | 0.0008978249 | 0.0002533103 | 0.374062797 | 0.1685887716 | 0.676193126 |
| **Gentamicin** | 0.1108027031 | 0.0490272876 | 0.431709250 | 0.1152119000 | 1.000000000 |
| **Ciprofloxacin** | 0.8385489076 | 0.0003633622 | NA | NA | 0.194295441 |
| **Co-amoxiclav** | 0.0857751742 | 0.0162110595 | 0.926173749 | 0.2554864522 | 0.005570945 |
| **Piperacillin-Tazobactam** | 0.0011324664 | 0.0014335714 | 0.000565947 | 0.6484983709 | 0.104526039 |
| **Meropenem** | 0.3025207858 | 0.3866889310 | 0.230473319 | 0.0563107490 | 0.064250082 |

**Supplementary table 2:** Paired T test to ascertain significant difference of mean OD600 between faecal and isolate pairs.

| Pair | P value |
| --- | --- |
| 1 | 0.011829607 |
| 2 | 0.002514331 |
| 3 | 0.045537335 |
| 4 | 0.183078015 |
| 5 | 0.022019967 |

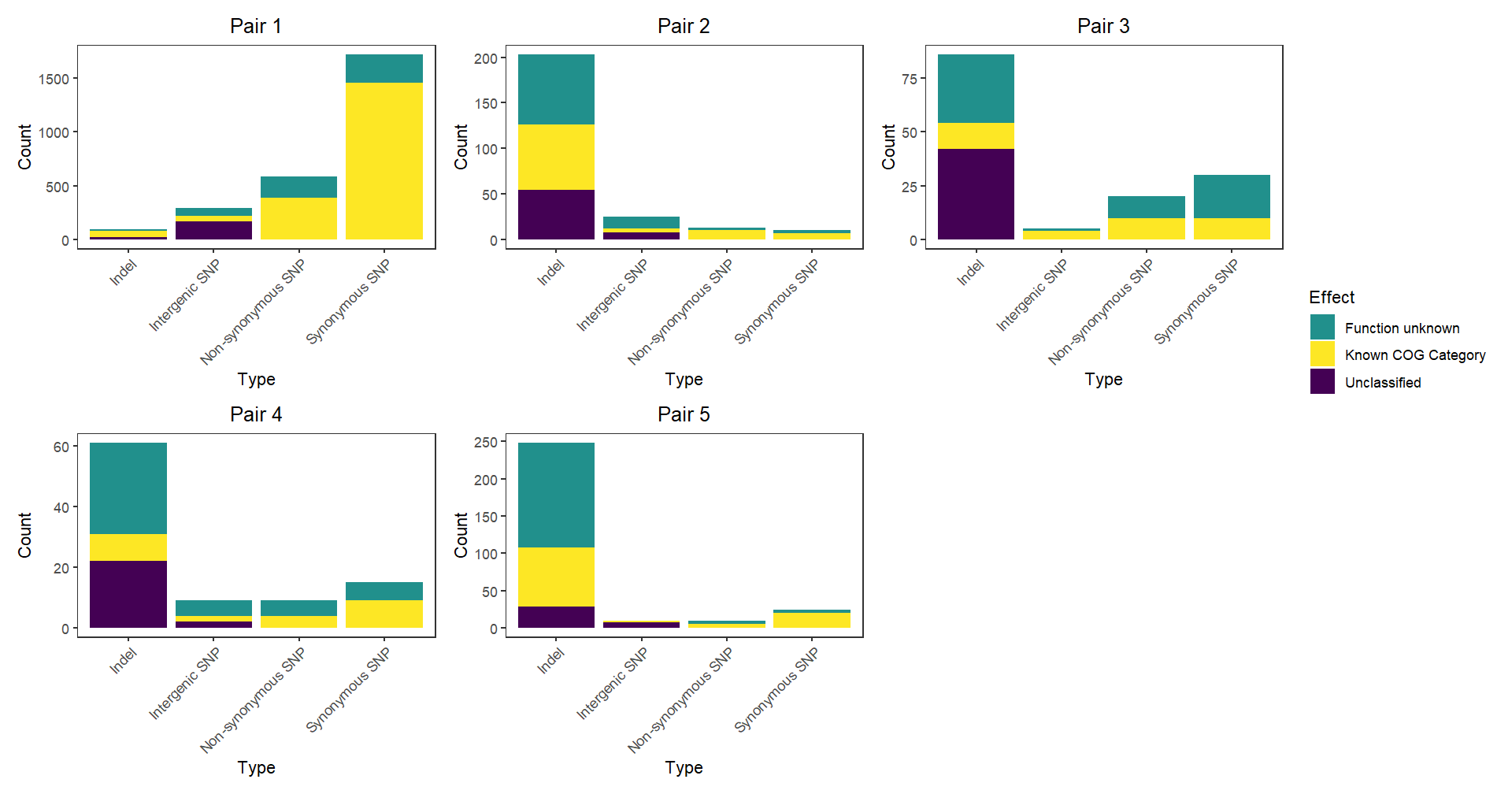

**Supplementary figure 3:** Mutation type in each isolate pair. On each chart, each bar represents the number of each type of mutation identified by Breseq (Indel, Intergenic SNP, Non-synonymous SNP, and Synonymous SNP) categorised by where mutation occurred. Function unknown/known COG category = mutation occurred in or within 100bp of a gene of either unknown function or known function respectively. Unclassified = mutation did not occur within 100bp of a gene.
